# Nested birth-death processes are competitive with parameter-heavy neural networks as time-dependent models of protein evolution

**DOI:** 10.64898/2026.02.02.702952

**Authors:** Annabel Large, Ian Holmes

## Abstract

Most statistical phylogenetics analyses use relatively simple continuous-time finite-state Markov models of point substitution to describe molecular evolution, often keeping sequence length fixed, ignoring insertions and deletions (indels) entirely, and making little (if any) allowance for variations in selection pressure due to interactions between amino acids. The simplistic assumptions of these models limit the realism of phylogenetics. We extend the TKF92 model - the canonical hierarchical model combining an outer birth-death process for indels with an inner finite-state Markov chain for substitutions - by introducing additional nesting and latent states, allowing for structural heterogeneity. We compare these TKF92 extensions (which are derived as exact solutions of instantaneous processes, and in which evolutionary time naturally appears as a matrix exponential coefficient) to two classes of neural seq2seq model that are not derived in such a way, but instead take evolutionary time as an input feature: the first class of model being constrained to enforce a TKF92-like structure, and the second lacking any such constraint. We evaluate the per-character perplexities of all models on splits of the PFam database of aligned protein domains. A nested TKF-based model with only 32,000 parameters is highly competitive with neural networks containing tens of millions of parameters, outperforming all but two of the neural architectures tested. Our results indicate that approaches grounded in molecular evolutionary theory may be more parameter-efficient and provide a better fit to real alignments than unconstrained alternatives, supporting the incorporation of CTMC-based model structure within future neural phylogenetic approaches.

## 1. Introduction

Substitutions and insertion-deletion (indel) events can alter protein function and drive complex evolutionary phenomena. Improved understanding of sequence evolution through time-dependent statistical models benefits areas such as epidemiology, biotechnology, and population genetics.

TKF91 and TKF92 are the canonical indel models for substitutions and indels, assuming an underlying instantaneous rate process (Thorne et al., 1991; 1992). These models take the form of hierarchically nested continuous-time Markov chains (CTMCs). At the outer level, a linear birth-death process over “links” describes changes in sequence length. At the inner level, each residue independently evolves under a finite-state CTMC. Exact finite-time solutions to these processes yield transition and emission parameters in a hidden Markov model (HMM) that defines the autoregressive likelihood P(*Z, Y*| *X, t*). Here *Z* is an alignment between ancestral sequence *X* and *Y*, assuming a specified evolutionary drift time *t*. Other related HMM-based indel models derive principled approximations to the underlying indel process (Löytynoja & Goldman, 2005; Redelings & Suchard, 2007; De Maio, 2021; Holmes, 2020).

In reality, each residue in a protein may experience different selection pressures due to biophysical constraints. Such rate variation can be accommodated in solvable CTMC-based models by composition, albeit in limited ways. For example, heterogeneity in substitution rate and mutation biases is routinely accounted for by introducing latent position-varying parameters into the model (Yang, 1994; Le & Gascuel, 2008; Prillo et al., 2024). Indel models are similarly amenable to such extensions and compositions (Holmes, 2004).

Use of these solvable CTMC-based models has both benefits and drawbacks. Benefits include directly interpretable rate parameters that quantify evolutionary dynamics, and that missing or artefactual data (e.g. sequence alignments) can be marginalized out of the likelihood. Drawbacks include that solvable CTMCs are limited in the complexity of the phenomena they can model. For example, indel rates are restricted by local sequence context and longer-range epistatic interactions, but incorporating this into a CTMC-based model is challenging.

An alternative to deriving HMM-based solutions is to use an autoregressive neural network to approximate the same empirical finite-time distribution P(*Z, Y*|*X, t*). Neural networks have gained traction in related areas of bioinformatics (Dotan et al., 2025; Becker & Stanke, 2024; Teng et al., 2024). Language models can capture evolutionary information from correlations in observed data (Hie et al., 2022; Brixi et al., 2025; Hayes et al., 2025) or by directly leverage phylogenetic information (Ye et al., 2025). EvoLSTM modeled pairwise alignments of DNA, but without evolutionary time (Lim & Blanchette, 2020). Combining neural sequence embeddings with molecular evolution models would increase expressive power, as neural architectures can capture complex multi-residue interactions, both local and distal, in ways that compositions of independent, solvable local CTMCs cannot (Wilburn & Eddy, 2020). However, neural parameters cannot be directly interpreted as quantifications of evolutionary dynamics. The Edit Flows model fits interpretable rates in text, although flow normalization methods maximize a different likelihood function than autoregressive methods (Havasi et al., 2025).

In this work, we explore several competing approaches to modeling molecular drift of sequences by substitution and indel events. All models provide an autoregressive likelihood of the form P(*Z, Y* | *X, t*). We curate a dataset of protein domain alignments from the PFam database. We compare several related HMM-based models (Thorne et al., 1991; 1992; Löytynoja & Goldman, 2005; Redelings & Suchard, 2007; Holmes, 2020) based on their cross-entropy on withheld alignments. TKF92 emerges as the best of the simple HMM-based models, so we extend TKF92 via mixtures and compositions at various levels of the hierarchically nested process. The resulting HMM-based elaborations of TKF92 gain expressivity while remaining exactly solvable. Our most expressive HMM is, to our knowledge, the first HMM-based indel model to allow indel rates to depend on local sequence context. We also develop two classes of neural transducers (Graves, 2012; Yeh et al., 2019). One neural model learns site- and sample-specific substitution and indel rates for a TKF92 model, yielding some interpretability. In contrast, the other uses generic neural network modules without any evolution-specific architectural components. Our innovation is to allow the alignment to explicitly guide cross-attention during neural network training. The motivation is that this may provide an inductive bias towards a Markovian evolutionary process suitable for phylogenetics. Finally, we compare the HMM-based elaborations against the neural methods based on their cross-entropy on the same withheld alignments as before. The models developed as solvable elaborations of TKF92 prove to be competitive with neural methods while using orders of magnitude fewer parameters.

All our proposed methods are constructed to satisfy what we call the “alignment-Markovian” property. The autoregressive likelihood P(*Z, Y*| *X, t*) is constrained to be strictly Markovian with respect to the alignment, though not necessarily with respect to the sequences themselves. HMM-based models have even stricter constraints and, thus, are alignment-Markovian by design. The main benefit of this property is that it allows the alignment to be marginalized out, as with TKF92 and prior HMM-based models.

After the initial deposit of this preprint, two independently developed and closely related works were deposited to bioRxiv and arXiv: PEINT (Koehl et al., 2026) and CoSiNE (Lu et al., 2026). Both address the problem of time-dependent deep learning models of protein sequence evolution, and we were pleased to find broad convergence in motivation across all three works. PEINT employs a seq2seq encoder-decoder architecture with evolutionary time provided as an explicit input feature, and leverages pretrained ESM embeddings to capture epistatic interactions. In contrast to our approach, cross-attention is not constrained by an alignment, evolutionary time does not enter through a matrix-exponential parameterization, and the model is not a neural transducer derived from an underlying instantaneous process. CoSiNE takes a more mechanistic direction that is complementary to ours: it parameterizes a CTMC with network-predicted, context-dependent substitution rates, and proves rigorously that this formulation constitutes a first-order approximation to a richer, sequence-wide substitution process. However, CoSiNE does not model insertions and deletions; its primary application is antibody affinity maturation, where point substitutions dominate. We conjecture that an analogous first-order approximation result may hold for our model in the joint substitution-and-indel setting, where the TKF structure plays a role analogous to CoSiNE’s CTMC; we regard this as an interesting direction for future theoretical work. We thank the authors of both papers for stimulating the field, and note that the present work was developed fully independently.

## 2. Background and prior work

### 2.1. Alignment-Markovian models

Let Ω be an alphabet of symbols (e.g. amino acids), and Ω_*g*_ = Ω∪ {*ϵ*} be this alphabet augmented with a gap symbol. A pairwise alignment (with all-gap columns excluded) may be considered a string over the alphabet of alignment columns, 𝒜= (Ω_*g*_×Ω_*g*_) {(*ϵ, ϵ*)}. Let *A* ∈𝒜 ^*^ be an alignment and let 𝒳: 𝒜^*^ → Ω^*^ and 𝒴: 𝒜^*^ →Ω^*^ be projection functions extracting, respectively, the unaligned ancestor and descendant sequences *X* = 𝒳 (*A*) and *Y* = 𝒴 (*A*).

The models we consider follow an *alignment-Markovian assumption*: the distribution of the next alignment column *A*_*k*_ is restricted to depend on the full ancestral sequence, the already-emitted portion of the descendant sequence, the lapsed evolutionary time, and the gap profile of the previous alignment column *A*_*k*−1_ (i.e. whether the ancestor token is a gap, the descendant token is a gap, or neither are gaps). This yields the factorization

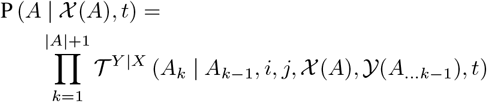

with *i* = |𝒳 (*A*_…*k*−1_) |and *j* = | 𝒴 (*A*_…*k*−1_) |.The term in the product is the *next-column conditional probability*

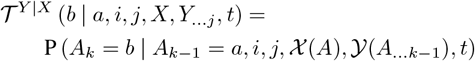

for the next alignment column *b* = *A*_*k*_ given the previous alignment column *a* = *A*_*k*−1_, current ancestor and descendant sequence positions (*i, j*), full ancestral sequence context *X* = 𝒳 (*A*), autoregressive descendant sequence context *Y*_…*j*_ =𝒴 (*A*_…*k*−1_), and lapsed evolutionary time *t*.

An alignment represents an evolutionary hypothesis, rarely observable directly. For given sequences *X, Y*, the most likely alignment can be inferred with the Viterbi algorithm, or alignment can be marginalized out of the likelihood altogether using the Forward algorithm (Durbin et al., 1998).

### 2.2. Pair hidden Markov models

The Pair HMM is a special case of an alignment-Markovian model where the next-column probability depends only on the previous alignment column and the next ancestral symbol to be aligned. We use the Moore machine framing of an HMM, where emissions are associated with states, so the next-column conditional probability factors into transition and emission probabilities

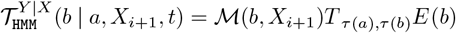

Here *T* denotes the time-dependent transition matrix, defined by the indel model; *E* (…) denotes the time-dependent emissions scoring function, defined by the substitution model and equilibrium distribution; ℳ (*b, X*_*i*+1_) is an indicator function that returns 1 if the given ancestor token is contained in the alignment column, and 0 otherwise; and *τ* (…) is the state type of the alignment column, i.e. its gap profile. The allowable state types are match `(M)` for columns containing two non-gap symbols (*x, y*); insert `(I)` for ancestral gaps aligned to descendant symbols (*ϵ, y*); delete `(D)` for ancestral symbols aligned to descendant gaps (*x, ϵ*); and sentinel tokens at sequence end, Start `(S)` and End (E).

### 2.3. F81 and GTR-LG08 point substitution models

TKF92-based models rely on point substitution models to describe evolution at individual sites of the sequence. We consider two forms for these substitution model: F81, a simple model parameterized by its equilibrium distribution (Felsenstein, 1981), and GTR-LG08, a more general, time-reversible model parameterized by an equilibrium distribution and a symmetric exchangeability matrix, which we fix to the LG08 exchangeabilities (Lanave et al., 1984; Le & Gascuel, 2008).

Let *s*(*t*) ∼ Subst(*Q, π*) denote the point substitution process, where *s*(*t*) is one residue at time *t, π* is the equilibrium distribution, and *Q* is the rate matrix (calculated from equilibrium distribution and exchangeabilities). The distribution of descendant symbols in match columns is P(*y* | *x, t*) = exp(*Qt*)_*xy*_, where exp(…) denotes the matrix exponential. The distribution of descendant symbols in insert columns is given by the equilibrium distribution.

With respect to the next-column conditional probability, the emissions scoring function *E* (*b*) returns the emission probability corresponding to the state type of column *b*.

### 2.4. TKF91 and TKF92 indel models: birth-death processes over links and fragments

A point substitution process alone cannot generate sequences with different lengths. TKF91 was the first indel model to describe both point mutations and single-residue indel events in a time-dependent manner (Thorne et al., 1991). However, under TKF91, estimates of indel event counts (and, hence, indel rates) tend to be inflated, since the model only permits one residue to be inserted or deleted per event. Consequently, alignments inferred under TKF91 tend to be unrealistic, with single-character indels scattered all over the sequence instead of being concentrated in a few multi-character indel events. The TKF92 model improves on TKF91 by allowing multiple residues to be inserted or deleted via a single event (Thorne et al., 1992). The nested structure of the TKF models is:

#### Outer model

The outer model is a linear, continuous-time birth-death process with immigration, where “mortal links” are born with insertion rate *λ* and die with deletion rate *µ*. There is also one “immortal link” that can never die. Each link is associated with a nested process, which evolves independently throughout the life of the link. Let *S*(*t*) ∼ Links(M; *λ, µ*) denote the hierarchically structured *links process*, where *S*(*t*) is a sequence at time *t* and M is the inner stochastic process acting at each mortal link.

#### TKF91 inner model

In TKF91, each mortal link represents a single residue evolving according to the finite-state CTMC

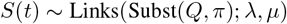

for some substitution rate matrix *Q* and equilibrium *π*.

#### TKF92 inner model

In TKF92, each mortal link represents a multi-residue fragment with independently evolving sites. The fragment *s*′(*t*) ∼ at time *t* is drawn from a *fragment process s*′(*t*) ∼ Frag(M; *r*) as follows. First a fragment length *K*∼ Geometric(*r*) (with *K* ≥ 1) is sampled. The stochastic process corresponding to each character in the fragment is then drawn from the substitution model M. The TKF92-distributed “sequence” of fragments is

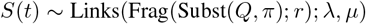

Both TKF91 and TKF92 are amenable to exact analysis, due to the tractability of the underlying birth-death model (Thorne et al., 1991; 1992). With respect to the next-column conditional probability, closed-form solutions to the TKF processes determine the transition probabilities between column state types, which populate the transition matrix *T*.

### 2.5. General Geometric Indel process

The TKF92 model was introduced by Thorne *et al* to remediate perceived problems with the TKF91 model: namely, its failure to model multi-character indel events. TKF92 solves this problem at the cost of introducing latent information into the sequence in the form of fragment boundaries. As a result, the underlying state space of TKF92 is not the set of sequences over the residue alphabet, but the set of sequences of multi-character fragments. Fragment boundaries are never observed, as they are technical conveniences that have no material interpretation. They must be imputed or (preferably) marginalized in any practical analysis.

This issue with TKF92 led several previous researchers to investigate CTMC models whose state consisted solely of a character sequence, with no hidden fragment structure. The canonical example is the General Geometric Indel (GGI) process (De Maio, 2021). This process assumes that indels occur at uniform random positions in the sequence and have geometrically distributed lengths, which corresponds to the maximum entropy assumptions for expected indel rates and lengths. The TKF92 model may be considered as an approximation to the GGI process; for the special case where indels are restricted to single residues, the GGI process reduces exactly to the TKF91 model.

Unlike TKF92 or TKF91, no closed-form solution is available for the finite-time transition probabilities of the GGI process. However, it is straightforward to simulate trajectories from the this process, and one can then compare the observed gap length distributions from these simulations with various approximations and heuristics. Several such approximations have been proposed and evaluated on simulated data (Knudsen & Miyamoto, 2003; Löytynoja & Goldman, 2005; Redelings & Suchard, 2007; De Maio, 2021; Holmes, 2020). Of these approaches, the renormalized ODE approach of (Holmes, 2020), which we refer to as H20, most accurately approximates the distributions of gap lengths observed in GGI-based simulations; it is closely followed by TKF92.

As part of our efforts to develop the theory of indel models beyond TKF92, we assessed these approaches not just as approximations to the idealized GGI process, but as explanatory models of real protein alignments. Along with H20, TKF91, and TKF92, we considered two other models, LG05 (Löytynoja & Goldman, 2005) and RS07 (Redelings & Suchard, 2007), both of which can be characterized as informed guesses at closed-form solutions.

### 2.6. Mixture of site classes

Using mixture distributions is the standard approach to modeling heterogeneous selection pressures across sites. (Yang, 1994; Le et al., 2008). Instead of being drawn from one point substitution process, each site is drawn from a categorical mixture of substitution processes.

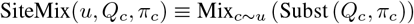

where Mix_*k*∼*p*_(*F* (*θ*_*k*_)) denotes a mixture of component models *F* (*θ*_*k*_) with component weights *p*_*k*_ and categorical class label *k*. Here, *c* ∼ *u* denotes site class.

By model design, the ancestral and descendant residues within an alignment column must share the same categorical label; site classes remain fixed and do not evolve. In order to evaluate alignment likelihoods, the unobserved latent class labels must be marginalized out. Conditioning on the ancestor sequence can make this marginalization cumbersome, so we instead evaluate the jointly-normalized likelihood P (*A*, 𝒳 (*A*) | *t*). The autoregressive decomposition uses the *next-column joint probability* 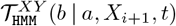 This likelihood has the same factorization as 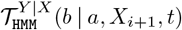 but the transition and emission probabilities include additional terms to account for the ancestral sequence.

## 3. Proposed models

### 3.1. TKF-based mixtures of processes

#### 3.1.1. Mixture of fragment classes

Instead of drawing from one fragment process, each TKF92 fragment is drawn from a categorical distribution of fragment processes. Each component fragment process also contains its own mixture over point substitution processes.

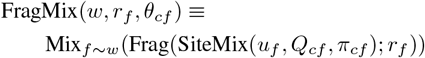

where *f* ∼ *w* is the fragment class label and *θ*_*cf*_ = (*u*_*f*_, *Q*_*cf*_, *π*_*cf*_) is the collection of parameters for the inner-level substitution processes. When evaluating the joint likelihood, latent site and fragment class labels appear as state indices in an expanded Pair HMM and can be marginalized out with the forward algorithm.

#### 3.1.2. Mixture of domain classes

Finally, we nest a mixture of TKF92-based models *inside* a TKF91 links model.

##### Outer model

The outer model is a TKF91 birth-death process where outer-level mortal links are born with insertion rate *λ*_0_ and deletion rate *µ*_0_.

##### Inner model

Each outer-level mortal link is associated with a subsequence of multi-residue fragments, which is generated by an inner-level TKF92 model. First, a TKF92 model is sampled from a categorical mixture; let *n* ∼ *v* be its domain class label. Next, an inner-level mortal link is either inserted with rate *λ*_*n*_ or deleted with rate *µ*_*n*_. The inner-level fragment length is then drawn from a mixture of fragment processes, and each site in the fragment is drawn from a mixture of substitution models

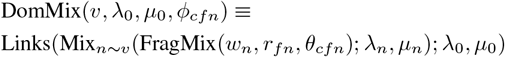

where *ϕ*_*cfn*_ = (*w*_*n*_, *r*_*fn*_, *θ*_*c,f,n*_, *λ*_*n*_, *µ*_*n*_) are the parameters for all lower-level mixture models. All latent classes manifest as additional states in the Pair HMM and are marginalized out of the joint likelihood with the forward algorithm.

### 3.2 Basic neural model

All previously described methods define an instantaneous evolutionary model and then attempt to solve the model’s finite-time transition probabilities (exactly or approximately). As an alternative, the *Basic neural model* is a kind of neural transducer that generates the aligned descendant sequence autoregressively, with the ancestral sequence and lapsed evolutionary time supplied as input features. We hypothesize that this model provides a reasonable approximation to the corresponding finite-time transition probabilities, despite the fact that this model is not based on a continuous-time Markov chain. Neural models do not contain latent class labels, so they are trained to maximize the conditional likelihood P (*A* | 𝒳 (*A*), *t*).

#### 3.2.1. Concise alphabet of alignment columns

There exist (| Ω| +1)^2^ possible alignment columns. However, only a subset of columns can have finite probability after conditioning on an ancestral sequence. Let Ω_aug_ = (Ω× {`M, I`}) ∪ {*ϵ*, `E`}be the concise alphabet that represents the possible columns after specifying an ancestral token. Let 𝒰 (*x*) be the |𝒜 ∪ {`E`}× |Ω_aug_| matrix that maps probabilities over the concise column alphabet to probabilities over the full column alphabet, conditioned on the ancestral token.

In the Basic neural model, the next-column conditional probability 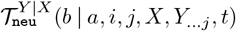 depends on position-specific embeddings from two neural networks:

- 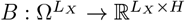 creates embeddings for an ancestral sequence of length *L*_*X*_, with embedding dimension *H*. Each embedding vector *B*(*X*)_*i*_ can be influenced by the entire ancestral sequence *X*.
- 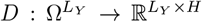 creates embeddings for a descendant sequence of length *L*_*Y*_. This embedding has causal context with respect to the descendant sequence: *D*(*Y*)_*j*_ ≡*D*(*Y* ⊗ *m*_≤*j*_)_*j*_ where *m*_≤*j*_ masks all positions after *j*.

The next-column conditional probability is generated by

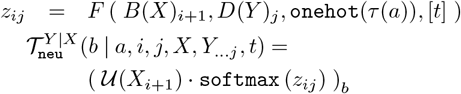

Where `layerNorm(…)` denotes layer normalization (Ba et al., 2016). Function *F* transforms the input features into logits *z*_*ij*_. Specifically, sequence embeddings at (*i*+1, *j*) are concatenated, layer normalized, and further concatenated with the previous column’s one-hot encoded state type and a single-element vector containing the lapsed evolutionary time. These column-wise embeddings are passed into a two-layer feedforward network with dropout, and the final logits are transformed into probabilities over the concise alphabet using softmax.

#### 3.2.3. Sequence embedding models

We evaluate three architectures for sequence embedding models *B* and *D*: a residual CNN (He et al., 2016; LeCun et al., 1989), an LSTM (Hochreiter & Schmidhuber, 1997) and a Transformer with rotary positional embeddings (RoPE) (Vaswani et al., 2017; Su et al., 2024). The residual CNN consists of a pre-norm residual block with a convolution layer, layer normalization, SiLU activation, and dropout (Ba et al., 2016; Elfwing et al., 2017; Srivastava et al., 2014). The Transformer follows a standard pre-norm architecture with a self-attention block and a two-layer multi-layer perceptron (MLP), each wrapped in residual connections, with layer normalization and SiLU activation.

### 3.3. Neural TKF model: a hybrid approach

We now combine instantaneous evolution processes with neural sequence embeddings. The *Neural TKF model* assumes a distinct TKF92+F81 model at every alignment column. The evolutionary model parameters are generated by neural networks from the available alignment-Markovian context. First, logits are generated by

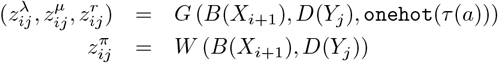

where *G* and *W* each transform their input features into logits 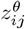, with *θ* = (*λ, µ, r, π*) being parameters of a TKF92+F81 model. Like before, sequence embeddings at (*i* + 1, *j*) are concatenated and layer normalized. *G* also concatenates the one-hot encoded state type of the previous column, but *W* omits this context. The resulting column-specific embeddings are passed into separate two-layer feedforward networks with dropout to generate logits 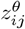. These logits are transformed into evolutionary model parameters by activation functions that ensure valid parameter values. These parameters yield site- and sample-specific TKF92+F81 transition and emission probabilities.

The Neural TKF next-column conditional probability is similar to that of the Pair HMMs.

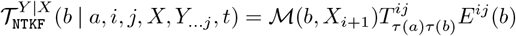

with a unique transition matrix *T* ^*ij*^ and emission scoring function *E*^*ij*^(*b*) for every position (*i, j*).

## 4. Methods

### 4.1. Dataset curation

Pairwise alignments are curated from Pfam 36.0 (Sonnhammer et al., 1997), specifically the Pfam-A seed alignments and phylogenetic trees. First, we discard: sequences repeated within an MSA (keeping only the first instance); sequences repeated across MSAs (i.e. across Pfams); peptides shorter than 10 amino acids, sequences containing nonstandard characters (B, O, U, X, Z); and Pfams with only one member. Twelve alignments were missing corresponding trees; these are imputed with FastTree 2.1.11 (Price et al., 2010), the same program used by Pfam (Finn et al., 2014).

We extract pairwise alignments from MSAs following the method described in (Prillo et al., 2023). Briefly, the two closest leaves in the tree are pruned and paired together. This process is repeated until the entire MSA is exhausted. If a single sequence remains unpaired, it is removed from the dataset. Sequence pairs are not realigned. Although observed alignments can never be assumed to be true, our models treat alignment as a latent variable that could be marginalized out. Since we assume time-reversible models, each pairwise alignment yields two samples: one with the first sequence as the ancestor and the second as the descendant, and another sample with the roles reversed. After postprocessing, less than 2% of pairwise alignments exceed 512 columns, and those that do have exceptionally many columns (up to 2,500). These longer alignments are removed to enable faster model prototyping.

In summary, we retained 19,909 Pfams and extracted 600,782 pairs of sequences to obtain a final training dataset of 1,201,564 pairwise alignments.

### 4.2. Train-dev-test partitions

Pfam clans and the remainder of families without clan labels are randomly assigned to one of ten splits. Once assigned, all the pairwise alignments of the given clan or family must be placed in the same split. This prevents homology information from unintentionally leaking to other splits. Assignment is balanced such that each split contains an approximately equal number of pairwise alignments. From these splits, three different permutations of train, dev, and test sets are created. Each partition has a 70:10:20 train-dev-test ratio.

### 4.3. Experiments

The optimization objective is to minimize the average of the conditional negative log-likelihood (NLL) over all alignments in training set 𝒟:

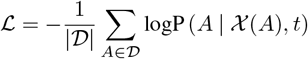

Training is repeated across three distinct train-dev-test partitions. Models are evaluated using the total NLL on the held-out test set, averaged over the three partitions. We also measure the exponentiated cross-entropy (ECE), which is computed by (i) normalizing each alignment’s NLL by the length of its descendant sequence, (ii) averaging these length-normalized values across all samples, and (iii) exponentiating the resulting average. This metric is essentially a per-column perplexity.

Our first investigation evaluates indel models that approximate or are special cases of the GGI process: TKF91, TKF92, LG05, RS07, and H20 (Thorne et al., 1991; 1992; Löytynoja & Goldman, 2005; Redelings & Suchard, 2007; Holmes, 2020). The equilibrium distribution is set to the observed amino acid frequencies, and the substitution probabilities are taken from the GTR-LG08 model. Branch lengths are provided by Pfam trees. We also briefly explored treating time as a latent variable and marginalizing over a grid of proposed times. This yielded similar likelihoods and conclusions at greater computational cost.

Moving forward, we use the observed amino acid frequencies as the equilibrium distribution only if there is one substitution process. Otherwise, distributions are fit during training. The F81 substitution model is used for emission probabilities. Our second investigation evaluates hierarchical mixtures of site, fragment, and domain classes and compares their Akaike information criterion (AIC) (Akaike, 1974) gains relative to the reference TKF92+F81 model. The number of mixture components is held constant across the lower hierarchical levels. For example, a model with two domain classes also includes two fragment classes and two site classes, yielding 2^3^ = 8 distinct combinations of labels. The numbers of mixture components tested are: 2-5, 10, 20, 30, 175, 500, and 900 for the mixture of site classes; 2-5, 10, 20, and 30 for the mixture of fragment classes; and 2-5 and 10 for the mixture of domain classes.

Our final investigation compares neural methods against the best mixture-based models. Initial trials tested six configurations of architectures formed by combining one of three sequence embedding architectures (Residual CNN, LSTM, and Transformer) with one of two prediction heads (Basic neural and Neural TKF). For each configuration, ancestor and descendant embedders share identical architectures and hyperparameters, except where necessary to enforce causal descendant sequence context.

We optimize a Transformer-based embedding architecture through hyperparameter sweeps while keeping the Neural TKF prediction head fixed. The sweeps vary the embedding dimension, number of Transformer blocks, number of self-attention heads, dropout rate, global learning rate, and weight decay. Performance is evaluated using the total NLL on a single dev set. The final Transformer-based model is called the “6-block” model, which reflects the optimal number of blocks found during the sweep. The initial Transformer configuration is called the “1-block” model. The same sequence embedding architecture is used for both the Neural TKF and Basic neural models, although only the former is explicitly optimized. We briefly explored alternative architectures for the Neural TKF prediction head (with the sequence embedders fixed), but the choice of sequence embedding architecture had more effect on model fit.

All models are trained using mini-batch stochastic gradient descent with the Adam optimizer (Kingma & Ba, 2017) on NVIDIA GeForce RTX 2080 Ti and RTX A6000 GPUs.

## 5. Results

### 5.1. Evaluation of approximations to the GGI process

Table 1 compares basic indel models and reports the total test set NLL, averaged over three train-dev-test partitions. Because TKF91 overestimates the number of indel events, it infers inflated indel rates and yields the poorest likelihoods. LG05 and RS07 make similar approximations to the finite-time transition probabilities. They have comparable performance and intermediate placement in the evaluation table, matching results from simulation studies. TKF92 demonstrates a slightly better fit to real alignments than H20, despite H20 proving better for simulated gap profiles.

**Table 1.** Comparing indel models by NLL of the held-out test set (lower NLL and ECE is better): “Total NLL (× 10^6^)” is the NLL of test set, summed over samples. “Average NLL” is the NLL averaged over samples. The best model, TKF92, is **bolded**.

| INDEL MODEL | TOTAL NLL<br>($\times 10^6$) | AVERAGE<br>NLL | ECE |
| --- | --- | --- | --- |
| **TKF92** | **71.58** | **280.240** | **6.1763** |
| H20 | 71.64 | 280.461 | 6.1875 |
| LG05 | 74.08 | 290.002 | 6.5310 |
| RS07 | 74.09 | 290.032 | 6.5321 |
| TKF91 | 75.70 | 296.362 | 6.7533 |

These initial investigations informed our subsequent model design and evaluations. Treating time as a latent variable and marginalizing it out increased computational cost with negligible affects to likelihood or indel model rankings. Therefore, we proceeded to evaluate likelihoods using Pfam branch lengths. Since TKF92 has a closed-form solution, is straightforward to elaborate further, and fits observed alignments best, it serves as the basis for our subsequent attempts to describe indel evolution more realistically. We accept the latent fragment boundaries inherent to TKF92 as a minor technical inconvenience that does not hurt performance.

### 5.2. Evaluation of TKF-based hierarchical mixtures

In Figure 1, the gain in AIC (Akaike, 1974) is plotted for all hierarchical mixture models (compared to TKF92+F81). This figure is generated from one train-dev-test partition, but trends are consistent across all three. Results demonstrate that increasing the number of site class components improves model fit, as compared to assuming only one substitution process; this is consistent with prior work (Yang, 1994; Le et al., 2008). However, across all latent class types, increasing the number of components yields diminishing improvements in model fit. In particular, as the number of components grows, models exhibit component collapse and redundancy. Better fit is achieved only by designing higher levels of process, as in our DomMix and FragMix models.

**Figure 1.**
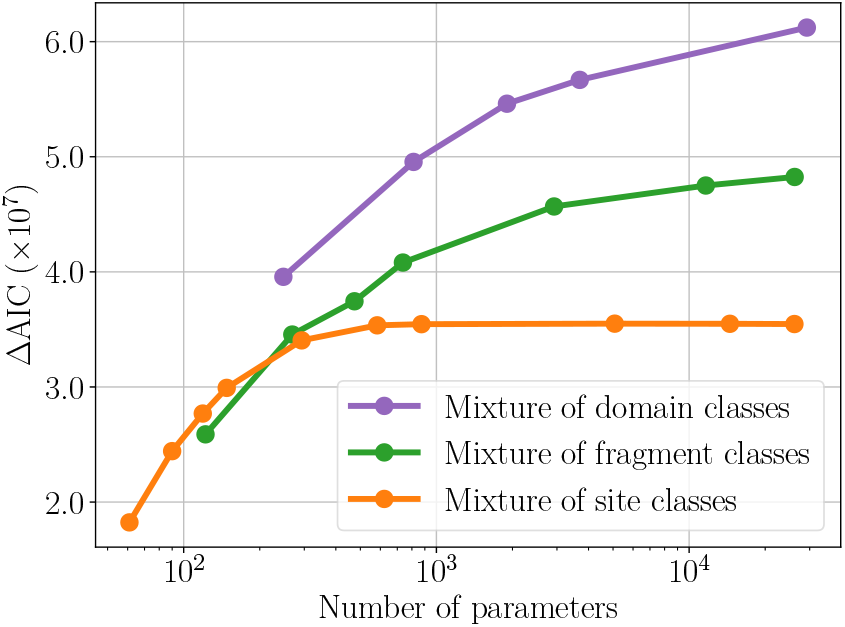
TKF-based mixtures of processes: gain in AIC over TKF92+F81 (higher is better): Each dot corresponds to a single trial with a fixed number of mixture components, detailed in Section 4.3. The x-axis shows the log-scaled number of model parameters and the y-axis shows gain in AIC over the reference model, TKF92+F81. Each curve asymptotically approaches (or is projected to approach) a limit on the performance gains obtained from increasing the number of components.

This behavior aligns with our expectations about how heterogeneous evolutionary forces are structured. Coordinating substitution mixtures within each fragment class reflects the expectation that residues collocated in sequence (and hence in space) are often subject to similar substitution pressures. Synchronizing mixtures of fragment processes (along with fragment-level indel rates) within each domain class reflects the fact that larger structural or functional regions have specific tolerances to indel events, in addition to known substitution biases. For example, the hydrophobic core of a globular protein tends to have poor tolerance for big insertions or hydrophilic amino acids.

Practically, optimizing a hierarchical mixture model requires few design choices. A relatively small number of parameters means no model ever overfits the training data. Although substantially larger mixtures would likely over-fit, increasing to such scale would be unjustifiable due to diminishing returns in performance.

### 5.3. Evaluation of neural and mixture-based models

Table 2 compares all neural models against the best-performing hierarchical mixtures and reports the total test set NLL, averaged over three train-dev-test partitions. The overall best-performing model is the Neural TKF with 6-block Transformer sequence embedders, followed by the same prediction head with LSTM embedders. However, by total NLL, the 10-component mixture of domain classes ranks third overall, despite using roughly three orders of magnitude fewer parameters than the neural methods. The difference in NLL between this mixture-based model and all neural models is smaller than the gap between LG05 and H20 (74.08 vs. 71.64 × 10^6^), which are trained on the exact same partitions and evaluated with identical branch lengths. The 30-component mixture of fragment classes also proves moderately comparable, as it outperforms the Basic neural model with 1-block Transformer sequence embedders.

**Table 2.** Comparing neural models against TKF-based hierarchical mixture models by NLL of held-out test set (lower NLL and ECE are better): “Number of parameters” is the number of trainable parameters in the model. Other column headings match those in Table 1. The best model, Neural TFK with 6-block Transformer sequence embedders, is **bolded**. We also mark the 10-component mixture of domain classes with a star (*⋆*) for its parameter efficiency and for achieving a lower NLL than all but two neural models.

| MODEL TYPE | MODEL SUBTYPE | NUMBER OF PARAMETERS | TOTAL NLL ($\times 10^6$) | AVERAGE NLL | ECE |
| --- | --- | --- | --- | --- | --- |
| **NEURAL TKF (SEC. 3.3)** | **6-BLOCK TRANSFORMER** | **43.55 M** | **61.72** | **246.867** | **5.5169** |
| NEURAL TKF | LSTM | 30.08 M | 61.85 | 247.388 | 5.5340 |
| NEURAL TKF | RESIDUAL CNN | 28.53 M | 62.47 | 249.864 | 5.6258 |
| NEURAL TKF | 1-BLOCK TRANSFORMER | 29.78 M | 62.99 | 251.962 | 5.6983 |
| BASIC NEURAL (SEC. 3.2) | LSTM | 27.00 M | 62.77 | 251.068 | 5.6877 |
| BASIC NEURAL | 6-BLOCK TRANSFORMER | 42.04 M | 62.82 | 251.282 | 5.6992 |
| BASIC NEURAL | RESIDUAL CNN | 26.47 M | 63.17 | 252.661 | 5.7455 |
| BASIC NEURAL | 1-BLOCK TRANSFORMER | 26.87 M | 64.04 | 256.156 | 5.8817 |
| MIXTURE OF |  |  |  |  |  |
| ... DOMAIN CLASSES (SEC. 3.1.2) | 10 COMPS. | ★ 29.23 K | ★ 62.14 | ★ 248.541 | ★ 5.7013 |
| ... FRAGMENT CLASSES (SEC. 3.1.1) | 30 COMPS. ($k_n = 1$) | 26.16 K | 63.87 | 255.479 | 5.9602 |
| ... SITE CLASSES (SEC. 2.6) | 30 COMPS. ($k_n = k_f = 1$) | 873 | 65.00 | 260.028 | 6.1238 |
| TKF92+F81 (SEC. 2.4) | N/A | 11 | 70.30 | 281.203 | 7.1398 |
| TKF91+F81 (SEC. 2.4) | N/A | 10 | 73.87 | 295.506 | 7.7942 |

When evaluated using ECE, all but two neural models achieve better fit than the 10-component mixture of domains model. That is, neural methods are, on average, more confident in their per-column predictions, whereas the mixture of domains model better captures the total likelihood. The discrepancy in model rankings suggests that performance may depend on underlying factors such as descendant length.

Finally, for any given sequence embedding architecture, the Neural TKF prediction head always achieved better model fit than the Basic neural prediction head; this is observed when calculating both total NLL and ECE. These results suggest that incorporating the inductive bias of evolutionary modeling improves neural network performance.

## 6. Discussion

In this work, we curate a training dataset of pairwise alignments and branch lengths from Pfam, and we re-evaluate indel models that approximate or reduce to the GGI indel process. (Holmes, 2020) demonstrated that H20 was better than TKF92 at recovering gap profile distributions, as simulated by the GGI process. However, our results show that TKF92, not H20, is a better fit for real data. This is surprising but not unexpected; simulated gap profiles are known to exhibit different length distributions than empirical ones (Wygoda et al., 2024).

We extend TKF92 into increasingly nested hierarchical mixture models and design two classes of neural models. Overall, the hierarchical mixtures were far easier to optimize than the neural methods. Our neural hyperparameter sweeps explored only a small fraction of the possible architectures. Additionally, the primary challenge during hyperparameter sweeps was scaling up without inducing overfitting to the training set. In contrast, the hierarchical mixtures had fewer design choices and never overfit.

During model evaluation, adding hierarchical levels gave a greater boost to model fit than increasing the number of components in any single level. Overall, hierarchical mixtures are competitive with neural methods at fitting Pfam alignments, despite having three orders of magnitude fewer parameters. Selection is heterogeneous across residues and context-dependent due to biophysical constraints. Neural models incorporate complex inter-residue interactions by design. Our results suggest that adding hierarchical mixtures to evolutionary models allows them to capture progressively more complex and realistic selection forces. In particular, the mixture of domains model introduces latent domain classes that contain coordinated fragment-level indel rates and substitution processes. Thus, surrounding sequence context can indirectly influence indel dynamics, despite the two being independent processes. This may explain why the performance of the mixture of domains model is comparable to much larger neural networks.

Finally, neural models with the inductive bias for evolutionary models consistently outperform their counterparts without bias. Similar conclusions have been drawn in several prior studies. (Prillo et al., 2024) found that fitting a point substitution model to every alignment column produced a variant effect prediction model that outperformed neural methods. (Hsu et al., 2022) also found that an augmented Potts model (which is suggested to capture evolutionary signals) was competitive with the neural protein language models of the time at predicting function from sequence.

As a counterpoint to noting the strong performance of neural models, we observe that an advantage of the HMM-based approaches, despite their inability to capture epistatic interactions between sites, lies in their amenability to exact statistical manipulation. As one illustration: the single-sequence HMM representing the stationary distribution of the mixture-of-domains nested TKF model (MixDom) can be distilled into a minimal order-1 HMM with 21×3 states — one for each element of {amino acids ∪ *BOS*}× {*S*, emit, *E*} — by converting expected amino acid adjacency statistics from the full MixDom HMM into transition probabilities of the reduced model, similarly to the approach described in (Holmes, 2020). A parallel construction yields a finite-state transducer that is order-1 with respect to both ancestor and descendant, with 21×21×5 states corresponding to {last ancestral residue}×{last descendant residue}× {*S, M, I, D, E*} ; its transition probabilities, when the transducer is composed with the distilled order-1 HMM, are proportional to expected usages of partially marginalized paths through the MixDom Pair HMM. Such an order-1 transducer can be composed naturally along the branches of a phylogenetic tree with the order-1 HMM at the root, yielding a full generative multi-sequence HMM (Holmes, 2003), and the likelihood of an observed MSA computed via a beam-search variant of the Forward algorithm — a workflow that integrates naturally with standard phylogenetic inference pipelines. Achieving analogous tractability with the neural seq2seq models would be considerably more challenging; one conceivable route would be a variational approach in which posteriors from a TKF-like model serve as a structured approximate family, but this remains to be developed. More broadly, the complementary strengths of mechanistic and neural approaches suggest many productive directions: richer latent-state structure in the HMM family, tighter integration of alignment uncertainty into neural training, and hybrid architectures in which a TKF-derived prior regularizes a neural likelihood are all natural extensions of the present work.

In conclusion, CTMC-based models remain a relevant framework for describing molecular evolution, even in the age of large neural networks.

## Acknowledgments

We thank Jeffrey Thorne, Yun Song, Sebastian Prillo, and Antoine Koehl for helpful discussions and feedback. This work was funded by NIH R01 grants HG004483, GM080203, and HG013117. GPUs were provided in part by the NVIDIA Academic Hardware Grant.

