## Appendix for "Nested birth-death processes are competitive with parameter-heavy neural networks as time-dependent models of protein evolution"

### A. Solutions to transition and emission models

For Pair HMMs, the next-column conditional probability takes the form

$$\mathcal{T}_{\texttt{HMM}}^{Y|X}(b \mid a, X_{i+1}, t) = \mathcal{M}(b, X_{i+1}) T_{\tau(a),\tau(b)} E(b)$$

The next-column joint probability $\mathcal{T}_{\texttt{HMM}}^{XY}$ has the same factorization. This section defines different forms of transition matrix $T$ and emissions scoring function $E$.

#### A.1. General time reversible model, and F81 as a simplification

Let $x \in \Omega$ be an ancestral symbol and $y \in \Omega$ be a descendant symbol. For the general time reversible (GTR) model, the substitution rate matrix, $Q$, is parameterized by a symmetric exchangeabilities matrix $\chi$ and the equilibrium distribution $\pi$. Off-diagonal elements are $q_{xy} = \chi_{xy}\pi_y$ Diagonal elements are set such that rows of $Q$ sum to zero. $Q$ is further normalized such that $-\sum_{x\in\Omega} \pi_x Q_{xx} = 1$. This calibrates time $t = 1$ to mean time for one substitution to occur.

The F81 substitution model is a special case of the GTR model, where all exchangeabilities are set to 1.

#### A.2. Emissions scoring function

Let $b = (x, y)$ be one alignment column, and let $\tau(b)$ return the column state type. The emissions scoring function for the next-column conditional probability returns $\mathrm{P}(y \mid x, t)$ for match columns, and $\mathrm{P}(y)$ at insert columns.

$$E^{Y|X}(b) = \begin{cases} \exp(Qt)_{xy} & \text{if } \tau(b) = \texttt{M} \\ \pi_y & \text{if } \tau(b) = \texttt{I} \\ 1 & \text{otherwise} \end{cases}$$

where exp denotes the matrix exponential.

The emissions scoring function for the next-column joint probability includes additional terms for the ancestral symbol. It evaluates $\mathrm{P}(x, y \mid t)$ at match columns, $\mathrm{P}(y)$ at insert sites, and $\mathrm{P}(x)$ at delete sites.

$$E^{XY}(b) = \begin{cases} \pi_x \exp(Qt)_{xy} & \text{if } \tau(b) = \texttt{M} \\ \pi_y & \text{if } \tau(b) = \texttt{I} \\ \pi_x & \text{if } \tau(b) = \texttt{D} \\ 1 & \text{otherwise} \end{cases}$$

#### A.3. TKF91

The finite-time transition probabilities of TKF91 come from solutions to the underlying birth-death process (Thorne et al., 1991). Let $\lambda$ and $\mu$ be the rate of inserting or deleting a mortal link, respectively. The probability $\alpha$ that a mortal link survives to time $t$, the insertion probability $\beta$, the orphaned insertion probability $\gamma$ and the stationary link-sequence extension probability $\kappa$ are

$$\begin{aligned} \alpha(\lambda, \mu, t) &= \exp(-\mu t) \\ \beta(\lambda, \mu, t) &= \frac{\lambda\left(\exp(-\lambda t) - \exp(-\mu t)\right)}{\mu \exp(-\lambda t) - \lambda \exp(-\mu t)} \\ \gamma(\lambda, \mu, t) &= 1 - \frac{\mu\beta}{\lambda(1-\alpha)} \\ \kappa(\lambda, \mu) &= \frac{\lambda}{\mu} \end{aligned}$$

The entries of the transition matrix give the probability of transitioning from one column state type to another $\mathrm{P}(\tau(b) \mid \tau(a), t)$.

For the next-column conditional probability, the TKF91 transition matrix is

$$T^{Y|X}_{\text{TKF91}} = \left(\begin{array}{c|ccccc} & \texttt{S} & \texttt{M} & \texttt{I} & \texttt{D} & \texttt{E} \\ \hline \texttt{S} & 0 & (1-\beta)\alpha & \beta & (1-\beta)(1-\alpha) & (1-\beta) \\ \texttt{M} & 0 & (1-\beta)\alpha & \beta & (1-\beta)(1-\alpha) & (1-\beta) \\ \texttt{I} & 0 & (1-\beta)\alpha & \beta & (1-\beta)(1-\alpha) & (1-\beta) \\ \texttt{D} & 0 & (1-\gamma)\alpha & \gamma & (1-\gamma)(1-\alpha) & (1-\gamma) \\ \texttt{E} & 0 & 0 & 0 & 0 & 0 \end{array}\right)$$

These terms appear in future transition matrices, so we abbreviate this matrix as $V = T^{Y|X}_{\text{TKF91}}$.

For the transition matrix used in the next-column joint probability, the rate of extending the ancestor sequence (i.e. with a match or a delete state) must be factored into the appropriate transitions.

$$T^{\text{XY}}_{\text{TKF91}} = \left(\begin{array}{c|ccccc} & \texttt{S} & \texttt{M} & \texttt{I} & \texttt{D} & \texttt{E} \\ \hline \texttt{S} & 0 & V_{\texttt{SM}} & V_{\texttt{SI}} & V_{\texttt{SD}} & V_{\texttt{SE}} \\ \texttt{M} & 0 & \kappa V_{\texttt{MM}} & V_{\texttt{IM}} & \kappa V_{\texttt{DM}} & (1-\kappa)V_{\texttt{ME}} \\ \texttt{I} & 0 & \kappa V_{\texttt{MI}} & V_{\texttt{II}} & \kappa V_{\texttt{DI}} & (1-\kappa)V_{\texttt{IE}} \\ \texttt{D} & 0 & \kappa V_{\texttt{MD}} & V_{\texttt{ID}} & \kappa V_{\texttt{DD}} & (1-\kappa)V_{\texttt{DE}} \\ \texttt{E} & 0 & 0 & 0 & 0 & 0 \end{array}\right)$$

#### A.4. TKF92

For the next-column conditional probability, the TKF92 transition matrix is

$$T^{Y|X}_{\text{TKF92}} = \left(\begin{array}{c|ccccc} & \texttt{S} & \texttt{M} & \texttt{I} & \texttt{D} & \texttt{E} \\ \hline \texttt{S} & 0 & V_{\texttt{SM}} & V_{\texttt{SI}} & V_{\texttt{SD}} & V_{\texttt{SE}} \\ \texttt{M} & 0 & \frac{1}{\nu}\left(r+(1-r)\kappa V_{\texttt{MM}}\right) & (1-r)V_{\texttt{MI}} & \frac{1}{\nu}(1-r)\kappa V_{\texttt{MD}} & V_{\texttt{ME}} \\ \texttt{I} & 0 & \frac{1}{\nu}(1-r)\kappa V_{\texttt{IM}} & r+(1-r)V_{\texttt{II}} & \frac{1}{\nu}(1-r)\kappa V_{\texttt{ID}} & V_{\texttt{IE}} \\ \texttt{D} & 0 & \frac{1}{\nu}(1-r)\kappa V_{\texttt{DM}} & (1-r)V_{\texttt{DI}} & \frac{1}{\nu}\left(r+(1-r)\kappa V_{\texttt{DD}}\right) & V_{\texttt{DE}} \\ \texttt{E} & 0 & 0 & 0 & 0 & 0 \end{array}\right)$$

where $\nu = r + (1-r)\frac{\lambda}{\mu}$. This form is slightly different from (Thorne et al., 1992), as it also conditions on the ancestral sequence consisting of fragments with unknown boundaries.

For the next-column joint probability, the transition matrix is

$$T^{\text{XY}}_{\text{TKF92}} = \left(\begin{array}{c|ccccc} & \texttt{S} & \texttt{M} & \texttt{I} & \texttt{D} & \texttt{E} \\ \hline \texttt{S} & 0 & \kappa V_{\texttt{SM}} & V_{\texttt{SI}} & \kappa V_{\texttt{SD}} & (1-\kappa)V_{\texttt{SE}} \\ \texttt{M} & 0 & r+(1-r)\kappa V_{\texttt{MM}} & (1-r)V_{\texttt{MI}} & (1-r)\kappa V_{\texttt{MD}} & (1-r)(1-\kappa)V_{\texttt{ME}} \\ \texttt{I} & 0 & (1-r)\kappa V_{\texttt{IM}} & r+(1-r)V_{\texttt{II}} & (1-r)\kappa V_{\texttt{ID}} & (1-r)(1-\kappa)V_{\texttt{IE}} \\ \texttt{D} & 0 & (1-r)\kappa V_{\texttt{DM}} & (1-r)V_{\texttt{DI}} & r+(1-r)\kappa V_{\texttt{DD}} & (1-r)(1-\kappa)V_{\texttt{DE}} \\ \texttt{E} & 0 & 0 & 0 & 0 & 0 \end{array}\right)$$

#### A.5. Emissions scoring function for mixture of site classes

With the mixture of site classes model, we marginalize over latent class labels $c \sim \text{Categorical}(u)$ inside the emissions scoring function.

$$E^{\text{XY}}_{\texttt{siteMix}}(b) = \begin{cases} \sum_c u_c \pi_{x,c} \exp(Q_c t)_{xy} & \text{if } \tau(b) = \texttt{M} \\ \sum_c u_c \pi_{yc} & \text{if } \tau(b) = \texttt{I} \\ \sum_c u_c \pi_{x,c} & \text{if } \tau(b) = \texttt{D} \\ 1 & \text{otherwise} \end{cases}$$

The next-column joint probability is evaluated as

$$\mathcal{T}^{XY}_{\texttt{HMM}}(b \mid a, X_{i+1}, t) = \mathcal{M}(b, X_{i+1}) T^{XY}_{\tau(a),\tau(b)} E^{\text{XY}}_{\texttt{siteMix}}(b)$$

#### A.6. Transition matrix for mixture of fragment classes

The set of Pair HMM states is expanded to include column states augmented with fragment class labels $f \sim \text{Categorical}(w)$. Let $b'$ and $a'$ be *state* containing the column's state and fragment label: $b' = (\tau(b), f)$. Start and end sentinel tokens do not have a fragment class. The new TKF92-based transition matrix is

$$T_{\texttt{fragMix}} = \begin{cases} r_f + (1-r_f)\, V_{\texttt{II}}\, w_g & \text{where } a' = (m,f), b' = (n,g), m = n = \texttt{I}, f = g \\ r_f + (1-r_f)\, \kappa\, V_{nn}\, w_g & \text{where } a' = (m,f), b' = (n,g), m = n \neq \texttt{I}, f = g \\ (1-r_f)\, V_{m\texttt{I}}\, w_g & \text{where } a' = (m,f), b' = (n,g), (m \neq n \text{ or } f \neq g), n = \texttt{I} \\ (1-r_f)\, \kappa\, V_{mn}\, w_g & \text{where } a' = (m,f), b' = (n,g), (m \neq n \text{ or } f \neq g), n \neq \texttt{I} \\ (1-r_f)\,(1-\kappa)\, V_{m\texttt{E}} & \text{where } a' = (m,f), \tau(b) = \texttt{E} \\ V_{\texttt{SI}}\, w_g & \text{where } \tau(a) = \texttt{S}, b' = (n,g), n = \texttt{I} \\ \kappa\, V_{\texttt{S}n}\, w_g & \text{where } \tau(a) = \texttt{S}, b' = (n,g), n \neq \texttt{I} \\ (1-\kappa)\, V_{\texttt{SE}} & \text{where } \tau(a) = \texttt{S}, \tau(b) = \texttt{E} \end{cases}$$

The fragment and site class labels are marginalized out with the Forward algorithm.

#### A.7. Transition matrix for mixture of domain classes

In the Pair HMM for the joint ancestral-descendant distribution, the $\texttt{S} \rightarrow \texttt{E}$ transition weight is $(1-\kappa)(1-\beta)$. Thus the probability that a $\texttt{M}$ state emits no sequence is $z_t = \sum_{n \in \mathcal{C}_n} v_n (1-\kappa_n)(1-\beta_n)$ where $\kappa_n$ and $\beta_n$ refer to birth-death parameters $\kappa$ and $\beta$ calculated from indel rates $\lambda_n$ and $\mu_n$.

The Single HMM for the stationary distribution over sequences represents a special case of this where $t = 0$ In this HMM, the start$\rightarrow$end transition weight is $1-\kappa$. Thus the probability that an $\texttt{I}$ or $\texttt{D}$ state generates a zero-length sequence is $\sum_{n \in \mathcal{C}_n} v_n (1-\kappa_n) \equiv z_0$.

Start with the TKF92 Pair HMM transition matrix and split the $\texttt{M}$, $\texttt{I}$, and $\texttt{D}$ states into non-emitting and emitting states. Lump the non-emitting $\texttt{M}$ and $\texttt{I}$ together as state $\texttt{A}$ and let the non-emitting $\texttt{D}$ be $\texttt{B}$.

This derivation builds off transition terms in the joint TKF91 transition matrix. For ease of notation, let $K = T^{Y|X}_{\text{TKF91}}$.

The $7 \times 7$ transition matrix for the exploded joint pair HMM is

$$v = \left( \begin{array}{c|ccccccc} & \texttt{S} & \texttt{M} & \texttt{I} & \texttt{D} & \texttt{E} & \texttt{A} & \texttt{B} \\ \hline \texttt{S} & 0 & (1-z_t)K_{\texttt{SM}} & (1-z_0)K_{\texttt{SI}} & (1-z_0)K_{\texttt{SD}} & K_{\texttt{SE}} & z_t K_{\texttt{SM}} + z_0 K_{\texttt{SI}} & z_0 K_{\texttt{SD}} \\ \texttt{M} & 0 & (1-z_t)K_{\texttt{MM}} & (1-z_0)K_{\texttt{MI}} & (1-z_0)K_{\texttt{MD}} & K_{\texttt{ME}} & z_t K_{\texttt{MM}} + z_0 K_{\texttt{MI}} & z_0 K_{\texttt{MD}} \\ \texttt{I} & 0 & (1-z_t)K_{\texttt{IM}} & (1-z_0)K_{\texttt{II}} & (1-z_0)K_{\texttt{ID}} & K_{\texttt{IE}} & z_t K_{\texttt{II}} + z_0 K_{\texttt{II}} & z_0 K_{\texttt{ID}} \\ \texttt{D} & 0 & (1-z_t)K_{\texttt{DM}} & (1-z_0)K_{\texttt{DI}} & (1-z_0)K_{\texttt{DD}} & K_{\texttt{DE}} & z_t K_{\texttt{DD}} + z_0 K_{\texttt{DI}} & z_0 K_{\texttt{DD}} \\ \texttt{E} & 0 & 0 & 0 & 0 & 0 & 0 & 0 \\ \texttt{A} & 0 & (1-z_t)K_{\texttt{MM}} & (1-z_0)K_{\texttt{MI}} & (1-z_0)K_{\texttt{MD}} & K_{\texttt{ME}} & z_t K_{\texttt{MM}} + z_0 K_{\texttt{MI}} & z_0 K_{\texttt{MD}} \\ \texttt{B} & 0 & (1-z_t)K_{\texttt{DD}} & (1-z_0)K_{\texttt{DI}} & (1-z_0)K_{\texttt{DD}} & K_{\texttt{DE}} & z_t K_{\texttt{DM}} + z_0 K_{\texttt{DI}} & z_0 K_{\texttt{DD}} \end{array} \right)$$

where $K \equiv K^{(0)}$ is the top-level TKF91 model with parameters $(\lambda_0, \mu_0)$. Let $U_{\Phi_1,\Phi_2}$ be the matrix formed from rows $m \in \Phi_1$ and columns $n \in \Phi_2$ of $v$. Consider the matrix of transitions between the empty states $\texttt{A}, \texttt{B}$

$$U_{\texttt{AB},\texttt{AB}} = \left( \begin{array}{c|cc} & \texttt{A} & \texttt{B} \\ \hline \texttt{A} & z_t K_{\texttt{MM}} + z_0 K_{\texttt{MI}} & z_0 K_{\texttt{MD}} \\ \texttt{B} & z_t K_{\texttt{DM}} + z_0 K_{\texttt{DI}} & z_0 K_{\texttt{DD}} \end{array} \right)$$

Summing over empty paths of all lengths (including zero-length) involving $\mathtt{A}, \mathtt{B}$

$$
\begin{aligned}
\sum_{k=0}^{\infty} U^k_{\mathtt{AB},\mathtt{AB}} &= (I - U_{\mathtt{AB},\mathtt{AB}})^{-1} \\
&= \frac{1}{\det(I - U_{\mathtt{AB},\mathtt{AB}})} \left( \begin{array}{c|cc} & \mathtt{A} & \mathtt{B} \\ \hline \mathtt{A} & 1 - z_0 K_{\mathtt{DD}} & z_0 K_{\mathtt{MD}} \\ \mathtt{B} & z_t K_{\mathtt{DM}} + z_0 K_{\mathtt{DI}} & 1 - z_t K_{\mathtt{MM}} - z_0 K_{\mathtt{MI}} \end{array} \right) \\
\det(I - U_{\mathtt{AB},\mathtt{AB}}) &= (1 - z_0 K_{\mathtt{DD}})(1 - z_t K_{\mathtt{MM}} - z_0 K_{\mathtt{MI}}) - (z_t K_{\mathtt{DM}} + z_0 K_{\mathtt{DI}}) z_0 K_{\mathtt{MD}}
\end{aligned}
$$

The effective nonempty $5 \times 5$ transition matrix (with $\mathtt{A}, \mathtt{B}$ summed out) is

$$
T' \equiv U_{\mathtt{SMIDE},\mathtt{SMIDE}} \; + \; U_{\mathtt{SMIDE},\mathtt{AB}} \cdot (I - U_{\mathtt{AB},\mathtt{AB}})^{-1} \cdot U_{\mathtt{AB},\mathtt{SMIDE}}
$$

The hierarchically nested Pair HMM has $2 + 5\,|\mathcal{C}_n|\,|\mathcal{C}_f|$ states: $\{\mathtt{SS}, \mathtt{EE}\} \cup \{\mathtt{UX}_{cf} : \mathtt{UX} \in \{\mathtt{MM}, \mathtt{MI}, \mathtt{MD}, \mathtt{II}, \mathtt{DD}\}, c \in \mathcal{C}_n, f \in \mathcal{C}_f\}$

The transition matrix $T_{\mathtt{DomMix}}$ for the mixture of domain classes model has entries

| Source $i$ ($\mathtt{UX}_{cf}$) | Destination $j$ ($\mathtt{VY}_{dg}$) | Terms in transition weight $T^{\text{DomMix}}_{ij} = K_{\mathtt{XE}} \times$ | $T'_{\mathtt{UV}}$ | $\times K_{\mathtt{SY}}$ | $+ \delta_{\mathtt{UV}} \delta_{cd} (\ldots + \delta_{\mathtt{XY}} \delta_{fg} (\ldots))$ |
|---|---|---|---|---|---|
| $\mathtt{SS}$ | $\mathtt{MY}_{dg}$ | | $T'_{\mathtt{SM}}$ | $v_d K^{(d)}_{\mathtt{SY}} w_{dg}$ | |
| | $\mathtt{II}_{dg}$ | | $T'_{\mathtt{SI}}$ | $v_d \kappa_d w_{dg}$ | |
| | $\mathtt{DD}_{dg}$ | | $T'_{\mathtt{SD}}$ | $v_d \kappa_d w_{dg}$ | |
| | $\mathtt{EE}$ | | $T'_{\mathtt{SE}}$ | | |
| $\mathtt{MX}_{cf}$ | $\mathtt{MY}_{dg}$ | $(1 - r_f) K^{(c)}_{\mathtt{XE}}$ | $T'_{\mathtt{MM}}$ | $v_d K^{(d)}_{\mathtt{SY}} w_{dg}$ | $+ \delta_{cd} \left( (1 - r_f) K^{(d)}_{\mathtt{XY}} w_{dg} + \delta_{\mathtt{XY}} \delta_{fg} r_g \right)$ |
| | $\mathtt{II}_{dg}$ | $(1 - r_f) K^{(c)}_{\mathtt{XE}}$ | $T'_{\mathtt{MI}}$ | $v_d \kappa_d w_{dg}$ | |
| | $\mathtt{DD}_{dg}$ | $(1 - r_f) K^{(c)}_{\mathtt{XE}}$ | $T'_{\mathtt{MD}}$ | $v_d \kappa_d w_{dg}$ | |
| | $\mathtt{EE}$ | $(1 - r_f) K^{(c)}_{\mathtt{XE}}$ | $T'_{\mathtt{ME}}$ | | |
| $\mathtt{II}_{cf}$ | $\mathtt{MY}_{dg}$ | $(1 - r_f)(1 - \kappa_c)$ | $T'_{\mathtt{IM}}$ | $v_d K^{(d)}_{\mathtt{SY}} w_{dg}$ | |
| | $\mathtt{II}_{dg}$ | $(1 - r_f)(1 - \kappa_c)$ | $T'_{\mathtt{II}}$ | $v_d \kappa_d w_{dg}$ | $+ \delta_{cd} ((1 - r_f) \kappa_d w_{dg} + \delta_{fg} r_g)$ |
| | $\mathtt{DD}_{dg}$ | $(1 - r_f)(1 - \kappa_c)$ | $T'_{\mathtt{ID}}$ | $v_d \kappa_d w_{dg}$ | |
| | $\mathtt{EE}$ | $(1 - r_f)(1 - \kappa_c)$ | $T'_{\mathtt{IE}}$ | | |
| $\mathtt{DD}_{cf}$ | $\mathtt{MY}_{dg}$ | $(1 - r_f)(1 - \kappa_c)$ | $T'_{\mathtt{DM}}$ | $v_d K^{(d)}_{\mathtt{SY}} w_{dg}$ | |
| | $\mathtt{II}_{dg}$ | $(1 - r_f)(1 - \kappa_c)$ | $T'_{\mathtt{DI}}$ | $v_d \kappa_d w_{dg}$ | |
| | $\mathtt{DD}_{dg}$ | $(1 - r_f)(1 - \kappa_c)$ | $T'_{\mathtt{DD}}$ | $v_d \kappa_d w_{dg}$ | $+ \delta_{cd} ((1 - r_f) \kappa_d w_{dg} + \delta_{fg} r_g)$ |
| | $\mathtt{EE}$ | $(1 - r_f)(1 - \kappa_c)$ | $T'_{\mathtt{DE}}$ | | |

where $K^{(n)}$ denotes an the inner-level, joint TKF91 with domain class label $n$ and indel rates $(\lambda_n, \mu_n)$.

### B. Neural network architectures

#### B.1. Sequence embedding architectures

Let $B(X)$ and $D(Y)$ be the neural networks that generate sequence embeddings with embedding size $H$. We will define $B(X)$ for concreteness, although the same formulae apply to $D(Y)$ (except where stated otherwise).

Both the ancestor and descendant sequences are first projected to a fixed-width vector space using a learned embedding matrices, $E$,

$$
h_0 = E \cdot \mathtt{onehot}(X)
$$

where $E \in \mathbb{R}^{H \times |\Omega|}$.

**Residual CNN** The Residual CNN (LeCun et al., 1989) uses convolutions to capture interactions between protein sites. This model follows a pre-layer normalization residual architecture with SiLU activation and dropout (Xiong et al., 2020; He et al., 2016; Elfwing et al., 2017; Srivastava et al., 2014).

$$
\begin{aligned}
h_1 &= \texttt{layerNorm}(h_0; \gamma, \beta) \\
h_2 &= \texttt{conv}(h_1; w_{\texttt{conv}}), b_{\texttt{conv}}) \\
h_3 &= \texttt{silu}(h_2) \\
h_4 &= \texttt{dropout}(h_3; p_{\texttt{B}}) \\
B_{\texttt{cnn}} &= h_4 + h_0
\end{aligned}
$$

The parameters are:

- $\gamma$, $\beta$: scale and bias for layer normalization
- $w_{\texttt{conv}} \in \mathbb{R}^{k \times H \times H}$: convolution kernel weights, with kernel width $k$
- $b_{\texttt{conv}} \in \mathbb{R}^{H}$: convolution kernel bias
- $p_{\texttt{B}}$: dropout rate

The ancestor embedding model centers the convolution kernel on the site of interest, giving it bidirectional context. The descendant embedding model uses a causal kernel, placing the site of interest at the rightmost position so its context only includes preceding sites.

We use an embedding size of $H = 1028$ and a dropout rate of $p_B = 0.20$. The ancestor embedding model uses a convolutional kernel width of 16, while the descendant model uses a kernel width of 8.

**LSTM** This refers to an RNN that uses LSTM units, as implemented by Flax function `flax.linen.OptimizedLSTMCell` (Hochreiter & Schmidhuber, 1997; Heek et al., 2024). The ancestor embedding model uses a bidirectional RNN, with forwards and backwards embeddings concatenated.

$$
\begin{aligned}
h_1 &= \texttt{LSTM}(h_0) \\
h_2 &= \texttt{reverse}(\texttt{LSTM}(\texttt{reverse}(h_0)) \\
B_{\texttt{LSTM}} &= h_1 \oplus h_2
\end{aligned}
$$

The descendant embedding model stops at $h_1$. We use a unidirectional RNN for embedding descendant sequences to maintain causal context. Thus, the descendant embedding dimension will be half that of the ancestor embedding dimension.

Again, we use an embedding size of $H = 1028$ and a dropout rate of $p = 0.20$.

**Transformer with Rotary Positional Encoding** Here, we use a pre-norm transformer block with standard dot-product self-attention and rotary positional encoding (Vaswani et al., 2017; Su et al., 2024), followed by a multilayer perceptron (MLP).

The self-attention block is

$$
\begin{aligned}
h_1 &= \texttt{layerNorm}(h_0; \gamma_1, \beta_2) \\
Q', K', V &= \texttt{linear}(h_1; w_{\texttt{Q,K,V}}, b_{\texttt{Q,K,V}}) \\
Q, K &= \texttt{RoPE}(Q', K') \\
h_2 &= \texttt{selfattention}(Q, K, V; w_{\texttt{O}}, b_{\texttt{O}}) \\
h_3 &= h_0 + \texttt{dropout}(h_2; p_B)
\end{aligned}
$$

Transformer parameters include $w_{\mathtt{Q,K,V}} \in \mathbb{R}^{H\times 3H}$, $w_{\mathtt{O}} \in \mathbb{R}^{H\times H}$, $b_{\mathtt{Q,K,V}} \in \mathbb{R}^{3H}$, and $b_{\mathtt{O}} \in \mathbb{R}^{H}$. The other parameters are the previously-described $\gamma_1$, $\beta_1$, and $p_{\mathtt{B}}$.

The MLP block is

$$\begin{aligned} h_4 &= \mathtt{layerNorm}(h_3; \gamma_2, \beta_2) \\ h_5 &= \mathtt{linear}(h_4; w_1, b_1) \\ h_6 &= \mathtt{silu}(h_5) \\ h_7 &= \mathtt{linear}(h_6; w_2, b_2) \\ B_{\mathtt{trans}} &= h_3 + \mathtt{dropout}(h_7; p_B) \end{aligned}$$

The parameters are $w_1, w_2 \in \mathbb{R}^{H\times H}$, biases $b_1, b_2 \in \mathbb{R}^{H}$, unique layer normalization parameters, $\gamma_2$ and $\beta_2$, and the same dropout rate as in the attention block, $p_{\mathtt{B}}$.

For the ancestor embedding $B_{\mathtt{trans}}$, the self-attention uses a padding mask. For the descendant embedding $D_{\mathtt{trans}}$, both a padding mask and a causal mask are applied.

For the 1-block Transformer, we use embedding size $H = 1,452$, dropout of $p_B = 0.20$, and two attention heads. For the final 6-block Transformer, we use a smaller embedding size $H = 756$ and 6 attention heads.

#### B.2. Architecture of Basic neural prediction head

The column-specific embeddings are created by concatenating the sequence embeddings, previous column's state, and evolutionary time. It will have embedding size $H'$.

$$h'_0 = \mathtt{layerNorm}\left(B(X)_{i+1} \oplus D(Y)_j; \gamma_F, \beta_F\right) \oplus \mathtt{onehot}(\tau(a)) \oplus [t]$$

where $h'_0 \in \mathbb{R}^{H'}$.

The prediction head, $F$, takes $h'_0$ as input and outputs logits used to calculate probabilities over the alignment-augmented alphabet, $\mathcal{T}^{Y|X}_{\mathtt{neu}}\left(b \mid a, i, j, X, Y_{\ldots j}\right)$

$$\begin{aligned} h'_1 &= \mathtt{linear}(h'_0; w_{\mathtt{F1}}, b_{\mathtt{F1}}) \\ h'_2 &= \mathtt{silu}(h'_1) \\ h'_3 &= \mathtt{dropout}(h'_2; p_{\mathtt{F}}) \\ z_{ij} &= \mathtt{linear}\left(h'_3; w_{\mathtt{F2}}, b_{\mathtt{F2}}\right) \\ \mathcal{T}^{Y|X}_{\mathtt{neu}}\left(b \mid a, i, j, X, Y_{\ldots j}\right) &= \left(\mathcal{U}(X_{i+1}) \cdot \mathtt{softmax}(z_{ij})\right)_b \end{aligned}$$

Parameters include dropout rates and layer normalization parameters specific to the prediction head, $p_{\mathtt{F}}$, $\gamma_F$, and $\beta_F$. The intermediate weights, $w_{\mathtt{F1}} \in \mathbb{R}^{H'\times H_{\text{int}}}$, project the concatenated features down to a smaller, intermediate size $H_{\mathtt{int}}$. The bias for the intermediate layer is $b_{\mathtt{F1}} \in \mathbb{R}^{H_{\text{int}}}$. The final projection weight, $w_{\mathtt{F2}} \in \mathbb{R}^{H_{\text{int}}\times|\Omega_{\text{aug}}|}$, and bias, $b_{\mathtt{F2}} \in \mathbb{R}^{|\Omega_{\text{aug}}|}$, transform the features into logits for each tuple and token in $\Omega_{\mathtt{aug}}$.

We use an internal embedding size $H_{\mathtt{int}} = 500$ and dropout rate $p_F = 0.20$

#### B.3. Architecture of Neural TKF prediction head

The site- and sample-specific TKF92+F81 parameters to derive are: $\pi_{ij}, \lambda_{ij}, \mu_{ij}$, and $r_{ij}$. Instead of calculating $\lambda_{ij}$ directly, we use an offset $o_{ij}$ such that $\lambda_{ij} = \mu_{ij}(1 - o_{ij})$. Constraining $0 \leq o < 1$ ensures $\mu > \lambda$.

Similar to before, we use a small feedforward network to post-process features after concatenation. We now abbreviate this as

$$\mathtt{FF}(x; w, b, p) = \mathtt{dropout}\left(\mathtt{silu}\left(\mathtt{linear}\left(x; w, b\right)\right); p\right)$$

Empirically, we find that site-specific rates can explode to extreme values during training. To account for this, we use a modified version of the sigmoid function,

$$\texttt{sigbound}(x, b_{\texttt{min}}, b_{\texttt{max}}) = b_{\texttt{min}} + \frac{b_{\texttt{max}} - b_{\texttt{min}}}{1 + \exp(-x)}$$

**Network $W$** produces logits for the equilibrium distribution, using the following feedforward network.

$$\begin{aligned} h'_{\pi,0} &= \texttt{layerNorm}\left(B(X)_{i+1} \oplus D(Y)_j; \gamma_\pi, \beta_\pi\right) \\ h'_{\pi,1} &= \texttt{FF}(h'_{\pi,0}; w_{\pi 1}, b_{\pi 1}, p_G) \\ z^\pi_{ij} &= \texttt{linear}\left(h'_{\pi,1}; w_{\pi 2}, b_{\pi 2}\right) \end{aligned}$$

where $\gamma_\pi, \gamma_\pi$ are unique scales and biases for layer normalization, $p_G$ is the dropout rate for the entire prediction head. Let the size of the embeddings after concatenation be $H'_e$. Then linear weights have sizes $w_{\pi 1} \in \mathbb{R}^{H'_e \times H_{\texttt{int}}}$, $w_{\pi 2} \in \mathbb{R}^{H_{\texttt{int}} \times |\Omega|}$. Bias vectors have sizes $b_{\pi 1} \in \mathbb{R}^{H_{\texttt{int}}}$ and $b_{\pi 2} \in \mathbb{R}^{|\Omega|}$.

Equilibrium distribution is obtained by applying the softmax function

$$\pi_{ij} = \texttt{softmax}\left(z^\pi_{ij}\right)$$

**Network $G$** produces logits for evolutionary model parameters associated with the indel process. It uses a similar set of layers as $W$, except column-specific embeddings also include the previous column's state, $\tau(a)$.

$$\begin{aligned} h'_0 &= \texttt{layerNorm}\left(B(X)_{i+1} \oplus D(Y)_j; \gamma, \beta\right) \oplus \tau(a) \\ h'_1 &= \texttt{FF}(h'_0; w_1, b_1, p_G) \\ z^o_{ij} &= \texttt{linear}\left(h'_1; w_{\texttt{o}}, b_{\texttt{o}}\right) \\ z^\mu_{ij} &= \texttt{linear}\left(h'_1; w_\mu, b_\mu\right) \\ z^r_{ij} &= \texttt{linear}\left(h'_1; w_{\texttt{r}}, b_{\texttt{r}}\right) \end{aligned}$$

where $\gamma$ and $\beta$ are unique scales and biases for layer normalization (for a total of three unique layer normalizations across the entire prediction head). With the new embedding size after concatenation, $H'_t$, the first set of linear weights and biases have sizes $w_1 \in \mathbb{R}^{H'_t \times H_{\texttt{int}}}$, $b_1 \in \mathbb{R}^{H_{\texttt{int}}}$. The final projection weights are sizes $w_{\texttt{o}}, w_\mu, w_{\texttt{r}} \in \mathbb{R}^{H_{\texttt{int}} \times 1}$, and the final biases $b_{\texttt{o}}, b_\mu, b_{\texttt{r}}$ are scalars.

For both $G$ and $W$, we use an internal embedding size $H_{\texttt{int}} = 500$ and dropout rate $p_G = 0.20$

Indel rates are obtained with the `sigbound` function, which constrains them to a fixed range.

$$\begin{aligned} o_{ij} &= \texttt{sigbound}\left(z^o_{ij}, o_{\texttt{min}}, o_{\texttt{max}}\right) \\ \mu_{ij} &= \texttt{sigbound}\left(z^\mu_{ij}, \mu_{\texttt{min}}, \mu_{\texttt{max}}\right) \end{aligned}$$

We use limits $o_{\texttt{min}} = 10^{-4}$, $o_{\texttt{max}} = 0.333$, $\mu_{\texttt{min}} = 10^{-4}$, and $\mu_{\texttt{max}} = 2$. Subsequently, $6 \times 10^{-4} \leq \lambda_{ij} \leq 1.9998$.

The TKF92 fragment length parameter is obtained with the standard sigmoid function.

$$r_{ij} = \texttt{sigmoid}\left(z^r_{ij}\right)$$

The emission function for the neural TKF model is a site-specific F81 model

$$E^{\text{ij}}(b) = \begin{cases} \exp(\rho_{ij} Q_{ij} t)_{xy} & \text{if } b = (x, y) \text{ where } x, y \in \Omega \\ (\pi_{ij})_y & \text{if } b = (\epsilon, y) \text{ where } y \in \Omega \\ 1 & \text{if } b = (x, \epsilon) \text{ where } x \in \Omega \\ 1 & \text{if } b = \texttt{E} \end{cases}$$

The transition matrix is also site-specific; it depends on the ancestral and descendant sequence indices $(i, j)$

$$T^{\text{ij}} = \left(\begin{array}{c|cccc} & \texttt{M} & \texttt{I} & \texttt{D} & \texttt{E} \\ \hline \texttt{S} & (1-\beta_{ij})\alpha_{ij} & \beta_{ij} & (1-\beta_{ij})(1-\alpha_{ij}) & 1-\beta_{ij} \\ \texttt{M} & \frac{1}{\nu_{ij}}\left(r_{ij}+(1-r_{ij})\kappa_{ij}(1-\beta_{ij})\alpha_{ij}\right) & (1-r_{ij})\beta_{ij} & \frac{1}{\nu_{ij}}(1-r_{ij})\kappa_{ij}(1-\beta_{ij})(1-\alpha_{ij}) & 1-\beta_{ij} \\ \texttt{I} & \frac{1}{\nu_{ij}}(1-r_{ij})\kappa_{ij}(1-\beta_{ij})\alpha_{ij} & r_{ij}+(1-r_{ij})\beta_{ij} & \frac{1}{\nu_{ij}}(1-r_{ij})\kappa_{ij}(1-\beta_{ij})(1-\alpha_{ij}) & 1-\beta_{ij} \\ \texttt{D} & \frac{1}{\nu_{ij}}(1-r_{ij})\kappa_{ij}(1-\gamma_{ij})\alpha_{ij} & (1-r_{ij})\gamma_{ij} & \frac{1}{\nu_{ij}}\left(r_{ij}+(1-r_{ij})\kappa_{ij}(1-\gamma_{ij})(1-\alpha_{ij})\right) & 1-\gamma_{ij} \end{array}\right)$$

Note that transitions with zero probability have been removed from this matrix.

#### B.4. Hyperparameter sweeps to optimize the Transformer sequence embedding models

We perform hyperparameter sweeps using the Transformer sequence embedders and the Neural TKF prediction head. This includes repeating the sequence embedding functions (with the number of repeats referred to as number of transformer blocks) and repeating the feedforward network $\texttt{FF}(\ldots)$ in the prediction head. The chosen values are marked with a star $(\star)$.

| **Parameter** | **Values tested** |
|---|---|
| Weight decay | $0.0, 10^{-4}, 10^{-3\star}$ |
| Learning rate | constant $10^{-4\star}$,<br>cosine decay with linear warm-up<br>(start at $10^{-4}$, peak at $10^{-3}$, end at $10^{-5}$) |
| Embedding size | $1450, 1028, 750^{\star}, 500, 625$ |
| Sequence embedding dropout rate, $p_B$ | $0.0, 0.2^{\star}, 0.3$ |
| Number of transformer blocks | $1, 2, 4, 6^{\star}, 10$ |
| Number of attention heads | $1, 2, 4, 6^{\star}, 10, 12$ |
| Prediction head dropout rate, $p_G$ | $0.0^{\star}, 0.1, 0.2$ |
| Number of times $\texttt{FF}(\ldots)$ is repeated in the prediction head | $1^{\star}, 5$ |
| Feedforward intermediate embedding dimension, in prediction head | $1500, 1000, 750, 500^{\star}, 250$ |
